# Detection of novel HIV-1 drug resistance mutations by support vector analysis of deep sequence data and experimental validation

**DOI:** 10.1101/804781

**Authors:** Mariano Avino, Emmanuel Ndashimye, Daniel J. Lizotte, Abayomi S. Olabode, Richard M. Gibson, Adam A. Meadows, Cissy M. Kityo, Eva Nabulime, Fred Kyeyune, Immaculate Nankya, Miguel E. Quiñones-Mateu, Eric J. Arts, Art F. Y. Poon

## Abstract

The global HIV-1 pandemic comprises many genetically divergent subtypes. Most of our understanding of drug resistance in HIV-1 derives from subtype B, which predominates in North America and western Europe. However, about 90% of the pandemic represents non-subtype B infections. Here, we use deep sequencing to analyze HIV-1 from infected individuals in Uganda who were either treatment-naïve or who experienced virologic failure on ART without the expected patterns of drug resistance. Our objective was to detect potentially novel associations between mutations in HIV-1 integrase and treatment outcomes in Uganda, where most infections are subtypes A or D. We retrieved a total of 380 archived plasma samples from patients at the Joint Clinical Research Centre (Kampala), of which 328 were integrase inhibitor-naïve and 52 were raltegravir (RAL)-based treatment failures. Next, we developed a bioinformatic pipeline for alignment and variant calling of the deep sequence data obtained from these samples from a MiSeq platform (Illumina). To detect associations between within-patient polymorphisms and treatment outcomes, we used a support vector machine (SVM) for feature selection with multiple imputation to account for partial reads and low quality base calls. Candidate point mutations of interest were experimentally introduced into the HIV-1 subtype B NL4-3 backbone to determine susceptibility to RAL in U87.CD4.CXCR4 cells. Finally, we carried out replication capacity experiments with wild-type and mutant viruses in TZM-bl cells in the presence and absence of RAL. Our analyses not only identified the known major mutation N155H and accessory mutations G163R and V151I, but also novel mutations I203M and I208L as most highly associated with RAL failure. The I203M and I208L mutations resulted in significantly decreased susceptibility to RAL (44.0-fold and 54.9-fold, respectively) compared to wild-type virus (EC_50_=0.32 nM), and may represent novel pathways of HIV-1 resistance to modern treatments.

**Author summary:** There are many different types of HIV-1 around the world. Most of the research on how HIV-1 can become resistant to drug treatment has focused on the type (B) that is the most common in high-income countries. However, about 90% of infections around the world are caused by a type other than B. We used next-generation sequencing to analyze samples of HIV-1 from patients in Uganda (mostly infected by types A and D) for whom drug treatment failed to work, and whose infections did not fit the classic pattern of adaptation based on B. Next, we used machine learning to detect mutations in these virus populations that could explain the treatment outcomes. Finally, we experimentally added two candidate mutations identified by our analysis to a laboratory strain of HIV-1 and confirmed that they conferred drug resistance to the virus. Our study reveals new pathways that other types of HIV-1 may use to evolve resistance to drugs that make up the current recommended treatment for newly diagnosed individuals.

## Introduction

There are currently six classes of antiretroviral drugs approved for treatment of HIV-1 infection, with protease inhibitors (PIs) and nucleoside and non-nucleoside reverse transcriptase inhibitors (NRTIs and NNRTIs) in most widespread use [1]. Integrase strand transfer inhibitors (INSTIs) are a more recent class of antiretroviral drugs targeting the virus-encoded integrase (IN) protein, which is responsible for inserting complementary DNA derived from the viral RNA genome into the genome of the host cell [2]. INSTIs are increasingly being used for individuals in low and middle-income countries (LMICs) for whom first- and second-line antiretroviral treatment (ART) regimens have failed, due to the emergence of drug resistance mutations (DRMs) to the PIs, NRTIs and/or NNRTIs that comprise these regimens [3]. With the exception of boosted PIs, there is typically a greater genetic barrier for HIV-1 to develop resistance to second-generation INSTIs, such as dolutegravir (DTG) and bictegravir (BIC), relative to other drugs [4, 5]. Even so, there are multiple well-characterized mutations conferring major and accessory resistance to INSTIs [6], where we employ the Stanford HIV Drug Resistance (HIVdb) guidelines for categorizing DRMs [7]. Most DRMs with major effects cause some level of cross-resistance to all drugs in this class — DTG, BIC, raltegravir (RAL) and elvitegravir (EVG) — with higher-level resistance to RAL and EVG compared to DTG and BIC.

Until recently, HIV-1 drug resistance studies have generally focused on individuals receiving ART in high income countries. The expansion of ART to over 18 million worldwide has turned attention to finding affordable methods for scaling up treatment monitoring and drug resistance testing. With the high volume of tests, some LMICs have adopted drug resistance genotyping by next generation sequencing (NGS) technologies [8, 9], in place of the more common but less scaleable Sanger sequencing approaches [10]. In addition to the ability to multiplex large numbers of patient samples into a single run [11], NGS has an added advantage of deep sequencing — where the same region of the virus genome is covered by sequences from hundreds or thousands of individual viruses in the sample — making it possible to reproducibly identify minority HIV-1 variants below the detection threshold of Sanger sequencing [12]. Despite their low frequencies within patients, these minority variants have clinical significance as they can anticipate the emergence of drug resistance and treatment failure [13]. Deep sequencing analysis of clinical HIV-1 samples in LMICs also provides a unique opportunity to identify potentially new HIV-1 polymorphisms associated with drug resistance in diverse HIV strains [14–16]. These opportunities also present significant bioinformatic challenges. Enormous amounts of sequence data must be processed accurately and efficiently, where sequencing error rates still exceed conventional Sanger methods [17]. The task of identifying novel associations between treatment outcomes and minority variants in diverse HIV-1 populations remains an open problem, and research has focused largely on HIV coreceptor tropism [18, 19] in populations predominantly affected by subtype B (but see [20]).

The global diversity of HIV-1 is structured into four phylogenetic groups, denoted by letters M-P [21]. The vast majority of infections worldwide are caused by group M viruses, which are further separated into subtypes that have distinct geographic distributions, possibly owing to early ‘founder effects’ in sub-Saharan Africa [22]. The majority of research on DRMs has historically been carried out on HIV-1 subtype B, owing in part to the predominance of this subtype in North America and western Europe — this subtype represents only about 10% of the global HIV-1 pandemic [23, 24]. Fortuitously, clinical outcomes on first- and second-line ART appear to be largely independent of HIV-1 subtype [25–27]. However, several studies (reviewed in [28]) have shown that non-subtype B infections can accumulate DRMs in response to treatment along mutational pathways that are distinct from subtype B. For example, a novel DRM in HIV-1 RT (V106M) has been reported to confer resistance to the NNRTI efavirenz (EFV) that is characteristic of HIV-1 subtype C [29]. More recently, investigators determined that the HIV-1 integrase mutation G118R confers a high level of resistance to RAL in the circulating recombinant form CRF02 AG, where the glycine is highly conserved across subtypes [30]. According to that study, G118R had only been previously observed in cell culture on exposure to another second-generation INSTI (MK-2048).

Historically, Uganda has had one of the highest burdens of HIV/AIDS in the world, with an estimated 1.3 million people living with HIV-1. The majority of infections in Uganda are caused by HIV-1 subtypes A and D, followed by A/D inter-subtype recombinants and subtype C [31]. With increasing access to ART, the transmission of DRMs is becoming increasingly common with an estimated 5% to 9% of treatment-naïve individuals carrying at least one primary DRM [32]. The majority of individuals starting ART in Uganda are prescribed a first-line regimen based on EFV, tenofovir (TDF) and a second NRTI, whereas almost no one had received the World Health Organization-recommended [33] INSTI + 2 NRTIs initial regimen that is more common for first-line therapy in higher income settings [34, 35]. DTG was recently introduced into new first line treatment regimens across sub-Saharan Africa, but treatment with any INSTI still represents less than 1% of active first line treatments in LMICs [36]. In most LMICs, INSTIs have generally been reserved for those requiring a third-line treatment regimen, with RAL-based regimens being quite successful in treatment-experienced individuals with multidrug-resistant HIV [37–39]. Several mutational pathways reducing susceptibility to RAL have been described, including the major DRMs T66K, Y143R, Q148H/K/R, and N155H in HIV-1 integrase [4, 40–42]. Notably, all of these studies were carried out in predominantly (*≥* 90%) HIV-1 subtype B cohorts or *in vitro* with a subtype B laboratory clone.

Here, we describe a bioinformatic approach to detect potential novel DRMs from NGS data sets that include all within-host polymorphisms above a frequency of 1%. We have applied this method to HIV NGS data from a cohort of individuals with HIV-1 non-subtype B infections in Uganda, of whom a subset had experienced treatment failure on RAL-containing salvage regimens. Additionally, we have experimentally verified the resistance effects of the novel DRMs predicted by our bioinformatic analysis by drug susceptibility assays *in vitro*, and characterized these mutations in structural models of HIV-1 integrase.

## Methods

### Data collection

The study samples were collected from the Center for AIDS Research Laboratory at the Joint Clinical Research Centre (JCRC) in Kampala, Uganda [34]. Written informed consent was provided by all study participants. Ethical approval was obtained from JCRC and University Hospitals Cleveland Medical Center/Case Western Reserve University Institutional Review Boards (EM-10-07 and 10-05-35). All investigations have been conducted according to the principles expressed in the Declaration of Helsinki. Patient samples were assigned to one of four categories based on treatment history and clinical outcome records in the JCRC database: treatment-naïve, first-line treatment failures, second-line treatment failures, and treatment failures on RAL-based salvage regimens (RAL failure). Treatment failure was defined by the presence of either a viral load above 1,000 copies/mL and/or a CD4 cell count below 250 cells/mm^3^ in the period following treatment initiation. Although current definitions of treatment failure tend to focus on viral load measurements, we retained the criterion based on CD4 cell counts for consistency with historical practice in this treatment population.

### RNA extraction and PCR amplification

For each sample, viral RNA was extracted from 200*μ*L of plasma using a QIAamp viral RNA Mini Kit (Qiagen) according to the manufacturers instructions. Reverse transcription of the full-length HIV-1 integrase (IN)-coding region from extracted viral RNA and amplification was performed with the sense primer RTA9F (5-TATGGGGAAAGACTCCTAAATTTA-3) and antisense primer 3Vif (5-AGCTAGTGTCCATTCATTG-3) using a Superscript III single RT-PCR system with Platinum Taq DNA polymerase kit (Thermo Fisher Scientific) as per the manufacturers instructions. The complementary DNA product was purified using a Quant-iT Picogreen dsDNA assay kit (Thermo Fisher Scientific) and quantified using a Qubit fluorometer (Thermo Fisher Scientific). The region encoding integrase was amplified in two parts by nested PCR using the following sets of primers: (1) sense primer INTF1B (5’-AGGTCTATCTGGCATGGGTACC -3’) and antisense primer INTR1B (5’-GATTGTAGGGAATTCCAAATTCCTGCT-3’); (2) sense primer INTF2B2 (5’-CAGGAATTTGGAATTCCCTACAATCCCC-3’) and antisense primer INFR2B4 (5’-TGTC TATAAAACCATCCCCTAGCTTTCCC-3’).

### Library preparation and deep sequencing

Two overlapping IN-PCR regions corresponding to the 288 amino acids of HIV-1 IN were sequenced with the MiSeq NGS platform (Illumina). The amplicons were purified with Agencourt AMPure XP (Beckman Coulter) and quantified using the Quant-iT Picogreen dsDNA assay kit (Thermo Fisher Scientific), prior to adding adapters using the Nextera XT sample prep kit (Illumina) with dual indexing for a maximum of 384 unique tags. The resulting libraries were quantified, normalized and pooled for paired-end sequencing (2*×*300 nt) on the Illumina MiSeq platform. Signal processing, base calling and structural variant analysis were performed with the MiSeq Reporter Software (version 2.6, Illumina). We deposited the unprocessed FASTQ data in the National Center for Biotechnology Information (NCBI) Short Read Archive (BioProject accession number PRJNA554675).

### Site directed mutagenesis of I203M and I208L

The I203M and I208L mutations were created in HIV INT gene from pREC-NFL (NL4-3) backbone using in-house site directed mutagenesis protocol. Briefly, the HIV-1 IN coding regions were amplified with the following primers: Vif 3 reverse 1 (5’-GTCCTGCTTGATATTCACACC-3’); INTREXT (5’-AATCCTCATCCTGTCTAC-3’); and INTFEXT1 (5’-AGAAGTAAACATAGTAACAGACTCACA-3’). The I203M mutation was created using sense primer 5’-gcaggggaaaga atagtagacATGatagcaacagacatacaaac-3’ and the antisense primer 5’-gtttgtatgtctgttgctatCATgtctactattctttcccctgc-3’); I208L was created using the sense primer 5’-gaatagtagacataatagcaacagacTTGcaaactaaagaattacaaaaa-3’ and antisense primer 5’-tttttgtaattctttagtttgCAAgtctgttgctattatgtctactatt c-3’. The presence of the mutation in the plasmid and the propagated virus was confirmed by PCR followed by Sanger sequencing.

### Cells and antiviral compounds

TZM-bl, U87.CD4.CXR4 and HEQ293T cell lines were obtained through the AIDS Research and Reference Reagent Program (Division of AIDS, National Institute of Allergy and Infectious Diseases, U.S.) [43]. All cell lines were maintained in Dulbecco modified Eagle medium (DMEM) supplemented with 10% fetal bovine serum (FBS) and 100 *μ*g/ml penicillin-streptomycin. In addition, U87.CD4.CXR4 cells were maintained in the presence of 300 *μ*g/ml G418 (an aminoglycoside antibiotic) and 1 *μ*g/ml puromycin (Invitrogen, Carlsbad, CA). All cell lines were sub-cultured every 3-4 days at 37*^◦^*C under 5% CO_2_. The TZM-bl cells contained reporter luciferase and *β*-galactocidase reporter genes that were activated by expression of HIV *tat*. DTG and RAL were provided by Gilead Sciences (Foster City, CA, USA).

### Construction of HIV INT chimeric viruses

HIV full length integrase PCR products were cloned into pREC NFL IN/URA3 vector and *Saccharomyces cerevisiae* MYA-906 cells (ATCC) using the yeast homologous recombination-gap repair system [44]. Following homologous recombination, plasmids were extracted from the yeast cells and transformed into electrocompetent *Escherichia coli* Stbl4 cells (Invitrogen). Plasmids were extracted using Qiagen miniprep kits and plasmid DNA was quantified using a Nan-oDrop spectrophotometer (Thermo Scientific). The presence of the mutation in the generated plasmid was confirmed by sequencing. Chimeric pREC NFL INT plasmids were co-transfected into HEK293T cells (3 *×* 10^4^ cells/well) along with the complementing plasmid pCMV cplt using Fugene 6 reagent (Promega, Madison, WI) as described previously [44]. Virus was then propagated on U87.CD4.CXCR4 cells as described [44].

### Drug susceptibility assay in TZM-bl cells

HIV susceptibility to DTG and RAL was determined using TZM-bl cells. Briefly, 20,000 cells per well were exposed to wildtype (WT), I203M or I208L HIV-1 in presence of 10-fold dilutions of DTG or RAL (100 *μ*M to 10*^−^*^7^ *μ*M) and DEAE-dextran (1mg/ml) in 96-well tissue culture plates (Corning). The amount of virus added to each well was normalised to a multiplicity of infection (MOI) of 0.01 based on the infectious titer. After 48 hr incubation at 37*^◦^*C and 5% CO_2_, the infectivity of viruses was quantified by staining the cells with X-gal as described previously [45] and then counting the cells using an ImmunoSpot reader. The fold changes in the effective concentrations for 50% inhibition (EC_50_) and standard errors of the mean (SEM) were calculated based on two sets of experiments, each performed in quadruplicate. Drug sensitivity curves were generated using nonlinear regression curve fitting features of GraphPad Prism 8.0 software (GraphPad Software, Inc., San Diego, CA). Drug resistance is presented as fold change in EC_50_ between WT and mutant viruses.

### Sequence analysis

We processed the FASTQ files generated by the Illumina MiSeq platform using a customized version of the MiCall pipeline (https://github.com/PoonLab/MiCall-Lite) [17]. First, the pipeline extracts the empirical *ϕ* X174 error rates from the ‘ErrorMetricsOut’ binary InterOp file associated with the MiSeq run, and then censors bases in the FASTQ files associated with problematic cycle-tile combinations with error rates exceeding a cutoff of 7.5%. Next, the program cutadapt (version 1.11) [46] was used to filter the FASTQ read data for Illumina adapter sequences. The pipeline subsequently used the alignment program Bowtie2 (version 2.2.6) [47] to map the paired-end read data to the full-length sequence encoding HIV-1 integrase of the HXB2 reference genome (Genbank accession K03455). This preliminary mapping stage was followed by the iterative re-mapping of reads from the original FASTQ files to new reference sequences, which were progressively updated with the plurality consensus of reads that were successfully mapped in the previous iteration [17]. A mapping quality score cutoff of *Q* = 20 was applied at this stage to filter ambiguously mapped reads. The primary outputs of the pipeline included the sample-specific nucleotide consensus sequence, coverage maps, and the matrix of amino acid frequencies in the coordinate system of the HXB2 reference; insertions relative to this reference coordinate system were written to a separate output file.

We used a Python script to filter the amino acid frequency matrices generated by the pipeline described above, using a minimum coverage threshold of at least 1,000 mapped reads per amino acid. First, any matrix corresponding to a pair of FASTQ files was discarded if the overall number of reads mapped to the sample-specific consensus sequence was below this threshold. Next, any individual amino acid position below this coverage threshold was coded as missing data in the remaining frequency matrices. Additionally, any sites with discordant amino acid frequencies within the overlapping region of the two amplicons — *i.e.*, where the frequency of an amino acid exceeded the threshold in one amplicon but not the other — was also coded as missing data. Next, the script converted the amino acid frequency data for each sample into a long binary vector that indicated whether each of the 20 amino acids was observed above a frequency threshold of 1% at every position in the integrase gene; these outputs are herein denoted the low-threshold (LT) data set. This dichotomization step was repeated at a frequency threshold at 20% to produce the high-threshold (HT) data set. Hence, the amino acid frequency data from the aligned short reads were encoded into two presence-absence matrices, each comprising 20 *×* 288 = 5, 760 variables (columns) and one row for every pair of FASTQ files. Similar dichotomization approaches (*e.g.*, sparse binary encoding [18]) have previously been used for feature selection analyses involving amino acid polymorphisms [18, 48]. All subsequent analyses were replicated across these two data sets.

### Data imputation

A substantial number of patient samples (Supplementary Table S1) were sequenced more than once on the MiSeq platform to take advantage of the large number of index combinations and sequencing yield of this instrument. In other words, the number of rows in the presence-absence matrix produced by the previous step was greater than the number of patient samples. To incorporate the entire data collection without unnecessarily and arbitrarily discarding or pooling repeated measurements, we randomly down-sampled redundant rows to obtain a reduced presence-absence matrix with one row per sample, and repeated this procedure to yield 10 replicate matrices. All subsequent analyses were replicated across these matrices.

The next stage of our analysis employed a support vector machine (SVM) classifier to identify putative associations between amino acid polymorphisms and RAL failure. Although extensions of SVMs have recently been developed to handle missing data [49], the prevailing approach is to use a generic method to impute missing values prior to the SVM analysis. We used multivariate imputation by chained equations as implemented in the *R* package *mice* [50]. Based on the overall proportion of missing observations in our data sets (3.2% for the LT data set and 2.9% for the HT data set), the recommended minimum number of imputations was 3 [51]. We decided to generate 5 imputed data sets for each of the 10 normalized data sets from the previous section. Further, we duplicated this approach for the LT and HT data sets for a total of 5 *×* 10 *×* 2 = 100 imputed data sets. To speed up the multivariate imputation, we used the *quickpred* variable selection procedure implemented in the *mice* package to filter the data for potentially significant predictors based on a simple correlation statistic. We excluded sites with an absolute correlation with the group labels below 50% and output the remaining variables to a preliminary predictor matrix. Each imputation was run for 20 iterations instead of the default 5 iterations, and convergence was visually assessed using the trace line plots of estimates against iteration numbers. To increase the robustness of results from the SVM analysis, we filtered amino acid features with non-zero weights (based on their incorporation into support vectors) by a minimum frequency of 80% across imputations, *i.e.*, at least 40 out of 50.

### SVM analysis

The preceding imputation step yielded 100 large presence-absence matrices encoding the observed HIV-1 integrase amino acid polymorphisms across baseline and treatment failure samples. We analyzed each imputed data matrix with a support vector machine (SVM) in which the samples were mapped to a high-dimensional feature space based on the presence or absence of amino acids, which was encoded by +1 and *−*1, respectively, as stipulated by the soft-margin linear SVM model [52]. Our samples were recategorized into two groups of outcomes (labels): samples from patients who experienced treatment failure on a RAL-based regimen (abbreviated to ‘RAL failure’); and samples from treatment-naïve patients and patients who experienced treatment failure on first- and second-line regimens without integrase inhibitors (‘RAL naïve’). An SVM attempts to locate the hyperplane, defined by a subset of data points (the support vectors), that most effectively separates the training data into the two groups. We performed an SVM analysis with the *svm.fs* function from the *R* package *penalizedSVM* [53] using an L1-norm penalty. Compared to an unpenalized SVM, this penalty function aggressively zeroes-out the coefficients associated with features that are less informative for classifying the data, and thereby provides a framework for feature selection [54]. To calibrate the *λ* tuning parameter of the SVM model, which controls the severity of penalizing data points that cross the margin of the hyperplane, we used a discrete grid search to determine the optimal *λ* with minimal misclassification error by 5-fold cross-validation [55]. After training and cross-validation, we generated the final SVM model for the entire data set using the optimized *λ* parameter. For each of the 100 data sets, we extracted the average feature weights and counts from the SVM results.

To corroborate the assignments of the most positive or negative feature weights to specific amino acids per treatment group, we calculated the odds ratios to quantify the statistical associations between the amino acid and outcomes (group labels). We calculated odds ratios and 95% confidence intervals by unconditional maximum likelihood estimation (Wald method) as implemented in the *R* package *epitools*, adding a fixed *n* = 0.5 value to every single cell of its contingency table to avoid a division by zero error (Haldane-Anscombe correction [56]). Subsequently, we averaged the results for each feature across all 100 data sets. Additional details on the sequence processing and SVM analysis are provided as Supplementary Text S1.

### Drug resistance prediction

To obtain resistance predictions from the sequence data from the IN coding region, we generated a consensus sequence at a polymorphism threshold of 20%, such that any position with two or more nucleotide frequencies above this threshold was encoded as an ambiguous base using the corresponding IUPAC symbol. Further, we censored all positions with fewer than 50 mapped reads as missing data – this threshold was less stringent than the minimum number of mapped reads (1000) required for positions to be carried over to the SVM analysis, because the objective of the latter was to detect associations with polymorphisms at a minimum frequency of 1%. Since a minimum of 10 mapped reads (1% *×* 1000) was required to be interpreted as a real polymorphism, the same number of mapped reads was required to influence the consensus sequence at a cutoff of 20% (50 reads *×* 20% = 10 reads). Drug resistance prediction scores on the resulting consensus sequences were obtained for RAL using the Stanford HIVdb algorithm version 8.4 (updated 2017-06-16) [7].

### Subtype and phylogenetic analysis

We used the same consensus sequences generated for drug resistance prediction to predict subtypes and reconstruct the phylogeny. We used the SCUEAL algorithm in HyPhy [57] to generate subtype classifications and detect inter- and intra-subtype recombination. Next, we excluded predicted recombinant sequences and sequences that were classified as circulating recombinant forms (CRFs), and generated a multiple sequence alignment from the remaining sequences using MUSCLE (version 3.8.425) [58]. This alignment also incorporated the HIV-1 reference sequences curated by the Los Alamos National Laboratory (LANL) HIV Sequence Database (http://www.hiv.lanl.gov) for subtypes A1, C, D and G, where this selection of references was based on subtyping results from this study population. The alignment was manually inspected and refined in AliView (version 1.19-beta-3) [59]. We used jModelTest (version 2.1.10) [60] to select the most effective nucleotide substitution model based on the Akaike Information Criterion (AIC). Finally, we used PhyML (version 20160207) [61] to reconstruct a phylogenetic tree by maximum likelihood under the AIC-selected model with the default bootstrap support analysis (1,000 replicates). The tree was visualized and manually annotated for subtypes in FigTree (version 1.4.2, A. Rambaut, http://tree.bio.ed.ac.uk/software/figtree).

### Structural analysis

The *α*-helix that connects the catalytic core domain (CCD) to the C-terminal domain (CTD) of the HIV-1 integrase protein is not resolved in the available DNA bound three-dimensional (3D) structure (PDB ID 5U1C). We therefore modelled the coordinates of the missing *α*-helix region (a total of 22 amino acids derived from PDB ID 1EX4) into one monomer (chain A) of the original structure (PDB ID 5U1C) using the program MODELLER (version 9.21) [62] to create an extended structure (PDB ID 5U1C extended). Using the extended structure as template, we built structural models of the I203M, I208L and combined I203M and I208L mutations respectively. The protein refinement program 3Drefine was used for energy minimization and optimization of all models [63]. Furthermore, we used the molecular structure visualization program PyMOL (version 1.7.2.1, http://pymol.org; Schrödinger, LLC) to visualize the sites where the I203M and I208L mutations are located on the 3D structure (PDB IDs 5U1C, 5U1C extended and 2B4J) of the HIV-1 integrase protein [64].

## Results

### Data collection

We obtained plasma samples for a total of 380 patients receiving treatment for HIV-1 infection at the Joint Clinical Research Center in Kampala, Uganda, for genotypic drug resistance testing by deep sequencing on the Illumina MiSeq platform. Of the 328 samples from INSTI-naïve individuals (RAL naïve), 85 samples were categorized as treatment-naïve, 127 as first-line treatment failures, and 116 as second-line treatment failures. The remaining 52 samples (14% of total) were obtained from individuals who had experienced treatment failures on raltegravir-based salvage regimens (RAL failure). From these samples, we generated a total of *n* = 524 paired FASTQ files with a mean of 83,185 reads per pair. Using the MiCall pipeline [17], which iteratively re-maps read data to update sample-specific reference sequences, we mapped an average of 81,189 reads (97.6% of the raw totals) to the HIV-1 IN coding region from each sample. This pipeline enforced a number of coverage and quality filtering criteria (see Methods), including a minimum requirement of 1,000 read coverage per amino acid position. Consequently, we discarded *n* = 7 FASTQ files due to insufficient numbers of reads that mapped to the HIV reference.

The *ϕ*X174 control error rates associated with the two MiSeq runs used for these sequencing experiments displayed the typical exponential decay with increasing cycle number, starting at a median of 0.44% (interquartile range, IQR: 0.32%, 0.96%) and ending at 9.2% (6.7%, 15.6%; Supplementary Figure S1). The overall median error rate was 1.5% (0.49%, 3.7%), which was consistent with previously reported error rates for this platform. Out of a total of 22,800 tile-cycle combinations, 2,095 and 3,360 combinations with an error rate exceeding 7.5% from the respective runs were excluded from further analysis. These combinations were concentrated in the last 100 cycles of the second reads (80.3% and 74.6%). The median sequence length after mapping with soft clips was 498 (IQR 325, 525) nt, indicating that the majority of the dropped base calls due to bad tile-cycles could be compensated by high quality base calls in the first reads at the paired-end read merger step of the pipeline.

A key challenge in detecting genetic associations in deep sequencing data from rapidly-evolving pathogens like HIV-1 is that polymorphisms can be observed at many, if not most, sites. Hypothetically, there exists a frequency threshold that optimally separates polymorphisms caused by sequencing errors from actual variants with potential clinical significance. Because it was not feasible to replicate all downstream analyses for an exhaustive sample of frequency thresholds, we proceeded with 1% (low threshold, LT) and 20% (high threshold, HT) to dichotomize the amino acid frequency data into binary presence/absence values. These values were chosen on the basis of prior information on the expected error rate for this sequencing platform [17] and the detection limit of Sanger sequencing [10, 12], respectively.

After excluding reads with low map quality scores and censoring low quality or discordant base calls, we obtained a mean coverage of 18,907 and 19,076 reads per amino acid site for LT and HT data sets, respectively (Supplementary Figure S2). We restricted our subsequent analyses to amino acid polymorphisms, excluding variation due to insertions, deletions and premature stop codons. On average, we observed 10^−4^ insertions and 8 *×* 10^−5^ deletions per nucleotide, which was within the expected range of indel error rates for this sequencing platform [65]. Ambiguous amino acid polymorphisms due to low base quality or incomplete coverage affected a small fraction of the data sets (3.2% and 2.9% for the LT and HT data, respectively). These ambiguities were encoded as missing data and handled through multiple imputation.

### Sequence subtyping

Subtyping analysis of the majority consensus sequences derived from the NGS samples confirmed that the majority of samples were assigned to HIV-1 subtypes A (*n* = 159, 49.7%) and D (*n* = 70, 21.3%) as expected for this study population in Uganda. An additional *n* = 28 (8.4%) samples were predicted to be A/D recombinants, and *n* = 16 (4.2%) samples were predicted to be sub-type C. The remaining samples were assigned to other inter-subtype recombinants or could not be confidently assigned to a known subtype or circulating recombinant form. We found no significant association between predicted HIV-1 subtypes and RAL-based treatment failure (Fisher’s exact test, *P* = 0.5). Variation among sequences was best explained by the general time reversible model of nucleotide substitution with invariant sites and a gamma distribution to model rate variation across sites (GTR+I+G). We reconstructed a maximum likelihood phylogeny under this model to verify the subtype predictions from SCUEAL relative to curated HIV-1 subtype reference sequences (Supplementary Figure S3), where sequences assigned to subtypes A, C and D comprised monophyletic clades with high bootstrap support values (*>*85%).

### Drug susceptibility by genotyping

Applying the Stanford HIVdb algorithm to the NGS consensus sequences to predict resistance to RAL, we confirmed that less than one-third of RAL-based treatment failures (shortened to ‘RAL failures’) manifested the classical RAL resistance pathways [34]. The complete breakdowns of resistance prediction scores by HIV-1 subtype in the RAL naïve and RAL failure groups are summarized in Figure 1. Out of 52 consensus sequences from RAL failure samples, only 14 samples (26.9%) were predicted to have high-level resistance to RAL (score *≥* 60). One of these samples (score 120) harbored the major RAL resistance mutation G140A (in 99.12% of reads) in combination with the accessory mutation E138K (99.1%). An additional 10 samples carried the major mutation N155H at frequencies between 94.2% and 99.7%, and two samples carried the major RAL resistance mutation T66K (at 93.8% and 62.7%, respectively). Finally, two samples with high RAL resistance scores in this group carried the major mutation Q148R in combination with accessory mutation G163R. This relative lack of expected mutational pathways in the RAL failure group, quantified by the low number of patient samples with resistance prediction scores in the susceptible to low-level resistance range (Figure 1), motivated a more comprehensive analysis of HIV-1 integrase polymorphisms in the deep sequencing data.

**Figure 1:**
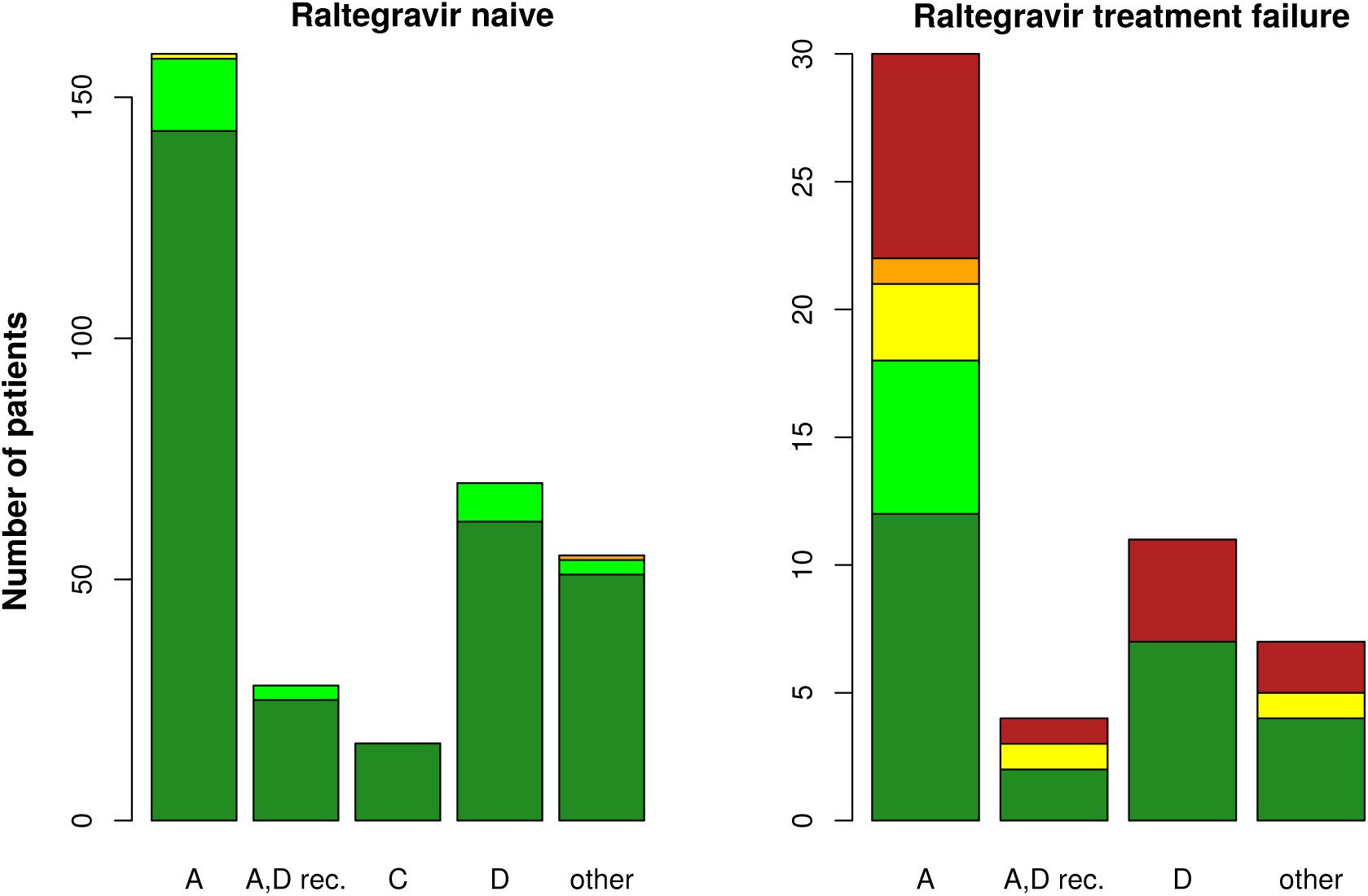
Distribution of RAL resistance predictions on sample consensus sequences by HIV sub-type and treatment outcomes. Resistance prediction scores were obtained by the Stanford HIVdb algorithm [7]. The study population was split into RAL naïve (left, *n* = 328) and RAL failure (right, *n* = 52) patients; accordingly, these plots are on different scales. Each stacked barplot stratifies patients by predicted HIV-1 subtype and categorizes the prediction scores into high-level (60+, red), intermediate (30-59, orange), low-level (15-29, yellow) and potential low-level resistance (10-14, light green), and RAL susceptible (below 10, dark green).

### Support vector machine analyses

A key challenge in detecting genetic associations in deep sequence data from rapidly-evolving infections like HIV-1 is that potentially any amino acid may appear at any position. We followed the sparse binary encoding approach in [18] so that every sample was represented by a total of 5,760 binary variables for 20 amino acids at 288 positions in the HIV-1 integrase reference. This is an unwieldy number of predictor variables for conventional association tests like logistic regression. Support vector machines (SVMs [66]) were developed to handle this sort of scenario, where the number of observed cases is vastly exceeded by the number of predictor variables, and have been employed in a number of studies of HIV variation [67, 68].

We used penalized SVMs to select features (polymorphisms) that most effectively separated patient samples into RAL naïve and RAL failure categories (labels; see Supplementary Text S1). Our results are summarized by the mean feature weights (the relative contributions of different polymorphisms to the separation of labels) for the low (1%) and high (20%) amino acid frequency threshold (LT and HT) data sets, respectively. Figures 2A (for LT) and 2B (for HT) display these results in comparison to the mean univariate odds ratios (ORs) for polymorphisms selected by the SVM analyses to provide more intuitive measures of association. To account for variation induced by missing data, we excluded features that were selected in fewer than 40 out of 50 imputed matrices for both LT and HT data sets, which disproportionately affected features with average weights close to zero. In general, we observed strong positive correlations between the SVM feature weights and mean ORs for the LT (Spearman’s *ρ* = 0.59, *P <* 2.2 *×* 10*^−^*^16^) and HT (*ρ* = 0.68, *P <* 2.2 *×* 10*^−^*^16^) data sets.

**Figure 2:**
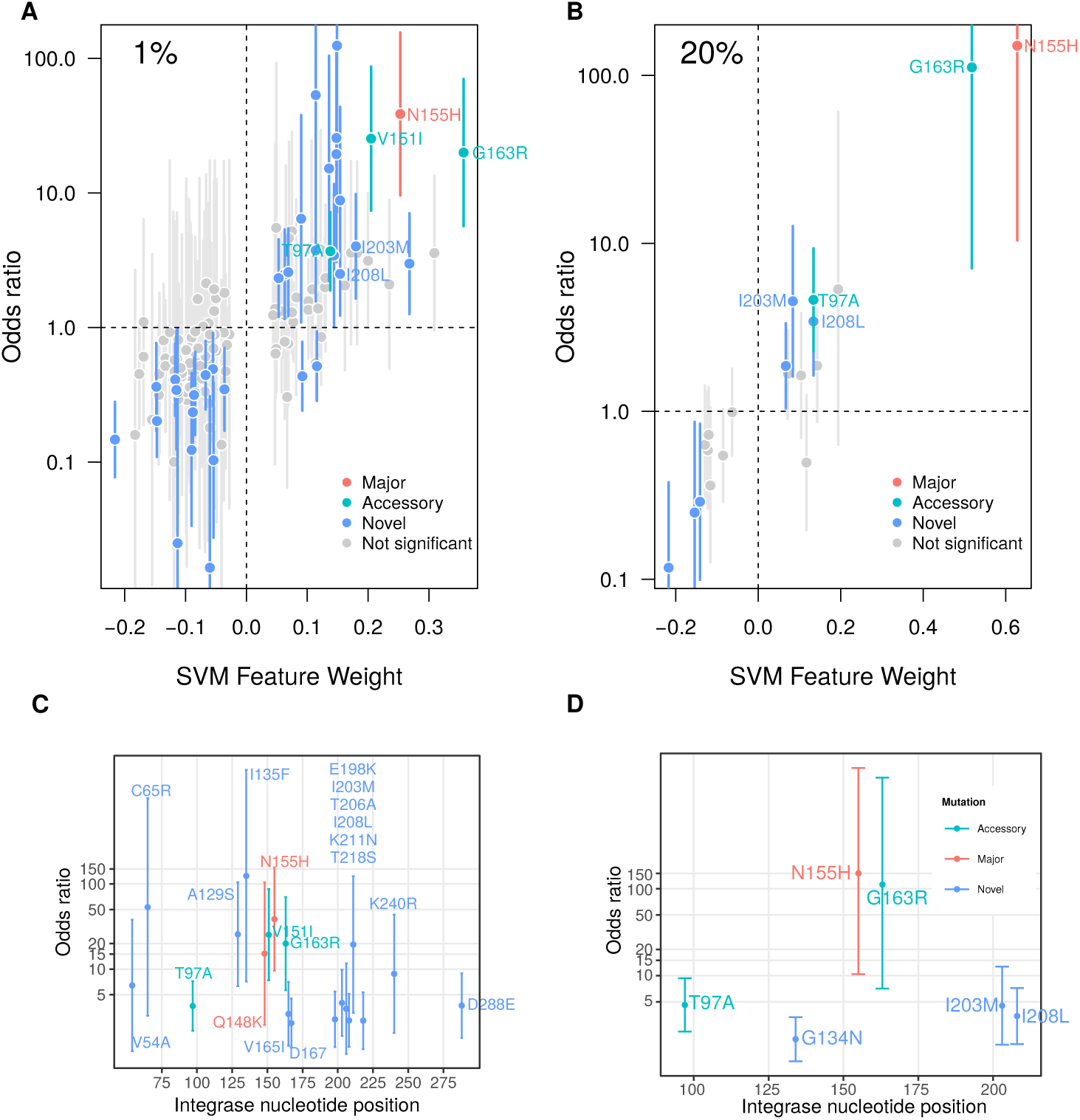
Summary of results from support vector machine analyses. (top) Feature weights and odds ratios for selected polymorphisms in the LT (A) and HT (B) databases. Each point corresponds to an amino acid polymorphism (feature) selected by support vectors. To reduce clutter and identify features robust to missing data, we removed all features selected in fewer than 40 out of 50 imputations. The *x*-axis corresponds to feature weights, and the *y*-axis represents the log-transformed mean odds ratio (OR) for each feature against the labels. Vertical lines indicate the empirical 95% confidence interval in ORs. Points are coloured grey if this interval spans 1 (not significant) and otherwise according to mutation categorization, if any, by the Stanford HIV Drug Resistance Database (see inset legend). (bottom) Positive weight selected from the SVM analysis and greater than one Odd ratio polymorphisms for the LT (C) and HT (D) data sets. The HIV-1 integrase reference coordinates are marked along the *x*-axis, and log-transformed OR mean and C.I. along the *y*-axis.

The entire set of amino acid polymorphisms selected by support vectors for the LT data set are summarized in Supplementary Table S2. To identify the most promising features from these results, we filtered the features that were selected in at least 40 imputations and where the mean lower 95% confidence limit in odds ratios was greater than 1. In total, we recorded 663 features selected by support vectors, of which 83 (12.5%) were reproducibly selected in at least 40 imputed matrices. Although the known major RAL resistance mutations T66K [69] and Q148R [70] had positive mean feature weights, they appeared in only 13 out of 50 imputed data sets (Supplementary Table S2). Only the major mutation N155H [71] was selected in a majority of imputations (all 50) with the fourth highest mean weight. An additional 8 features were known accessory or minor RAL resistance mutations (T66A [72], L74M [73], T97A [4], V151I [74], N155D [75], E157Q [76], G163R [74]) and R263K [77]); with the exception of R263K, all were assigned positive weights (Supplementary Table S2). Only G163R, V151I and T97A were selected in a majority (*>*80%) of analyses, and in fact were selected for all 50 imputed matrices. Overall, our filtering criteria selected the following polymorphisms in descending order of weight: G163R, V165I, N155H, V151I, I203M, T97A, K211N, A129S, D288E, K240R, Q148K, I135F, C65R, I208L and T218S. The relative locations of these polymorphisms are summarized in Figure 2C. For instance, we observed a cluster of 6 selected polymorphisms within the interval IN 198 to 218, which is distal to both the integrase active site and RAL binding site.

The higher frequency threshold (*<*20%) for the HT data set substantially reduced the effective number of amino acid polymorphisms; as a result, there were fewer features selected in the support vectors (*n* = 299; Supplementary Table S3). Applying the same criteria as above to identify the most significant features yielded the following six substitutions (in decreasing order of weight): N155H, G163R, T97A, I208L, I203M and G134N (Figure 2D). Of these features, only the primary mutation N155H and secondary mutations T97A and G163R have been previously described.

### I203M and I208L confer resistance to INSTIs *in vitro*

Based on these SVM analyses, we selected the novel substitutions I203M and I208L for further investigation, as they appeared in both lists of the most significant features from the LT and HT data sets. Figure 3 contrasts the intra-host frequencies of I203M and I208L in the context of known major RAL mutations, as defined by the Stanford HIV Drug Resistance Database. These frequency distributions revealed four distinct clusters of mutations among *n* = 30 (58%) patients failing a RAL-based treatment regimen and carrying at least one of these polymorphisms. Two clusters of patients in this RAL failure group carried one of two known major RAL resistance mutations Q148K/R or N155H. There were no patient samples containing the polymorphism Q148H. A third cluster comprised samples carrying the mutation I208L, and a fourth carried I203M and/or I208L to the exclusion of all known major RAL resistance mutations with the exception of Q148K/R. This pattern suggests that I203M and I208L may comprise novel mutational pathways for RAL resistance.

**Figure 3:**
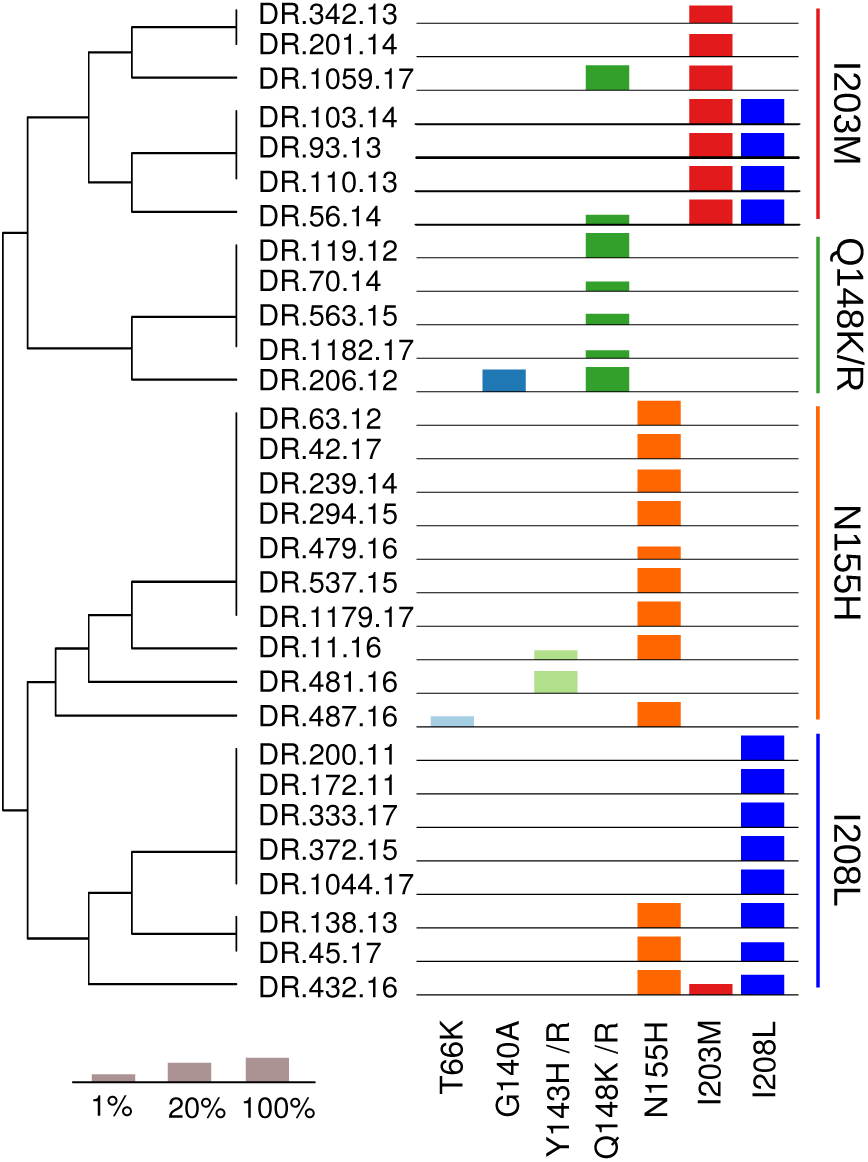
Summary of intra-host frequencies for known major RAL mutations and two candidate mutations I203M and I208L. Samples from *n* = 30 RAL failures patients carrying at least one of these mutations are each represented by a set of barplots representing mutation frequencies on the right. The height of each bar is proportional to 1 + log_10_(*p*) for *p* ranging from 1% to 100% (see legend in bottom-left). The vertical ordering of samples was determined by a hierarchical clustering analysis, with the corresponding dendrogram displayed on the left. Vertical bars have been added on the left to highlight the subsequent specific pathways.

To experimentally validate the predicted effects on resistance to RAL and other INSTIs, the I203M and I208L were first introduced into the IN coding-region of the NL4-3 HIV-1 proviral genome. The mutated proviral DNA was then transfected into 293T cells and the resulting virus was propagated on U87.CD4.CXCR4 cells *in vitro*. We then performed drug susceptibility assays on TZMbl cells exposed to the wild type and mutated HIV-1 and treated with varying concentrations of RAL and DTG. These experimental results are summarized in Figure 4. We confirmed that the presence of either I203M or I208L significantly decreased susceptibility to RAL (44.0-fold and 54.9-fold, respectively) compared to wild-type virus (EC_50_ = 0.32 nM). Furthermore, we found that I203M significantly decreased susceptibility to DTG (111-fold) and that there was no significant effect of the I208L IN (3.6-fold).

**Figure 4:**
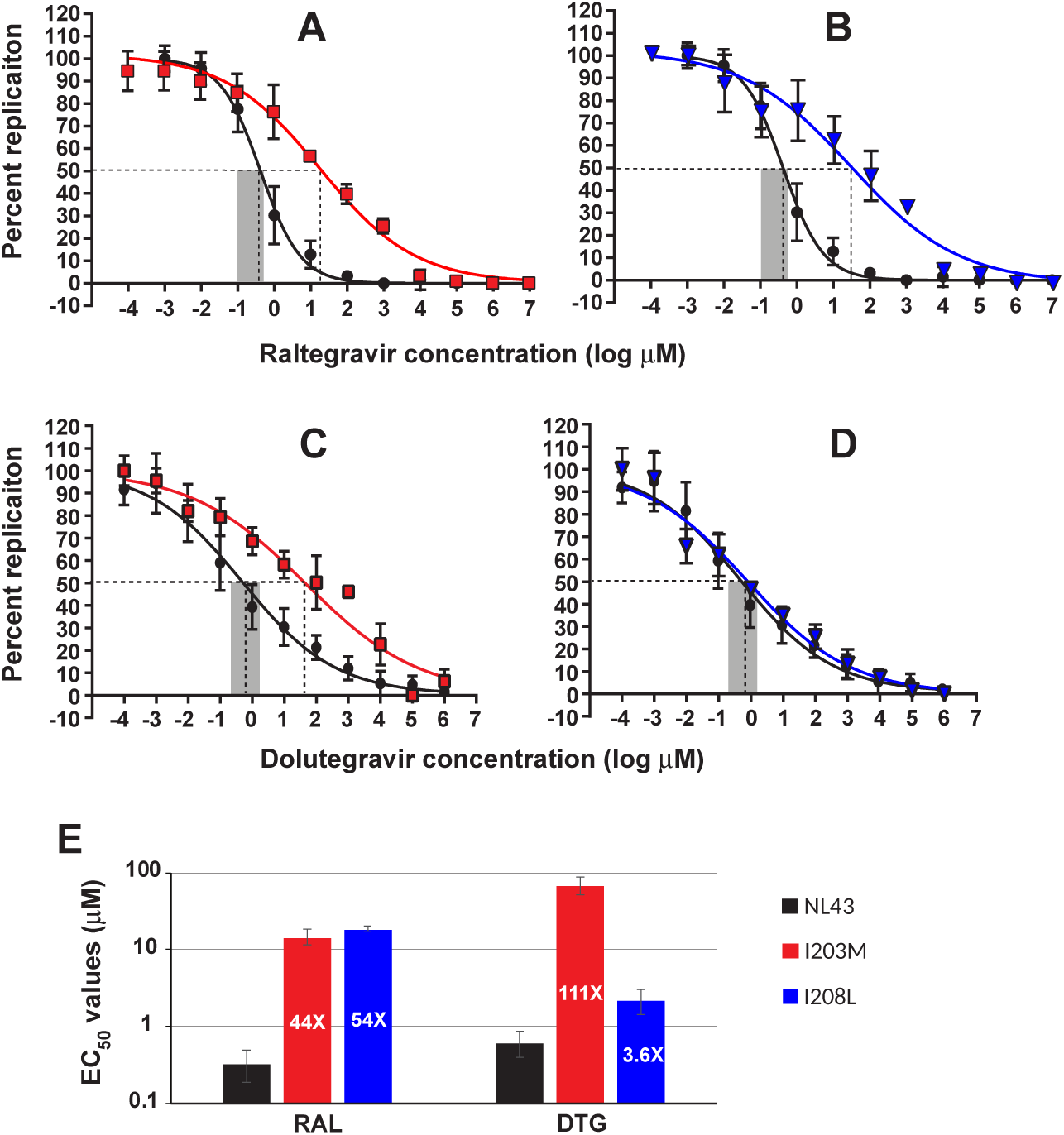
Drug susceptibility assays confirm the effects of I203M and I208L on INSTI resistance. The top row of plots correspond to raltegravir drug susceptibility curves for NL4-3 (wildtype, in black) and I203M (red, A) and I208L (blue, B), respectively. Points represent the mean and the error bars represent the standard error for a minimum of four replicates. Similarly, the middle row of plots correspond to the dolutegravir drug susceptibility curves for NL4-3 and I203M (C) and I208L (D). The barplot (E) summarizes EC_50_ fold change measurements for raltegravir (RAL) and dolutegravir (DTG) stratified by NL4-3 (black), I203M (in red) and I208L (in blue). Each point for curves in A-D corresponds to the mean and standard error of at least four replicate assays.

### I203M and I208L are natural polymorphisms

To assess whether I203M and I208L mutations are present as natural polymorphisms in the general population, we queried the Stanford HIV Drug Resistance Database (HIVdb; last accessed February 21, 2019) for all available HIV-1 IN sequences (*n* = 17, 605) from INSTI naïve and experienced patients and evaluated the frequencies of these variants in subtypes A, B and D (Table 2). This database comprised *n* = 1, 248 records from RAL-experienced individuals, of which only *n* = 129 were infected with a non-B subtype — in comparison, our study would contribute an additional *n* = 52 sequences, about 40% of the current number. The frequencies of I203M and I208L in the RAL-naïve category in the HIVdb database were 5.9% and 5.0%, respectively, suggesting these two mutations are natural polymorphisms that circulate at low frequencies (*>* 0.5%). We noticed only slight but statistically significant increases in these frequencies in association with the RAL-experienced category in HIVdb (Table 2). Although we observed more substantial increases in association with the RAL failure category in our study population (Table 1), this is not directly comparable to the RAL experienced category in the HIVdb database that comprises an unknown proportion of treatment failures. In addition, we also collected the frequencies of the known major RAL resistance mutations in HIVdb. These mutations were almost always observed with INSTI-experienced patients, being almost absent in INSTI naïve patients (Table 2).

**Table 1:** Frequencies of I203M and I208L polymorphisms in HIV-1 IN stratified by treatment for this study cohort and by minor allele frequency, *i.e.*, low threshold = 1% (LT) and high threshold = 20% (HT). Fisher’s exact tests (experienced vs. naïve): * = *P <* 0.05; ** = *P <* 0.01; *** = *P <* 0.001.

| Mutations | Low threshold |  |  | High threshold |  |  |
| --- | --- | --- | --- | --- | --- | --- |
|  | naïve | experienced |  | naïve | experienced |  |
| Total | 328 | 52 |  | 328 | 52 |  |
| I203M | 14 (4.3%) | 8 (15.4%) | ** | 8 (2.4%) | 6 (11.5%) | ** |
| I208L | 39 (11.9%) | 12 (23.1%) |  | 28 (8.5%) | 12 (23.1%) | * |

**Table 2:**
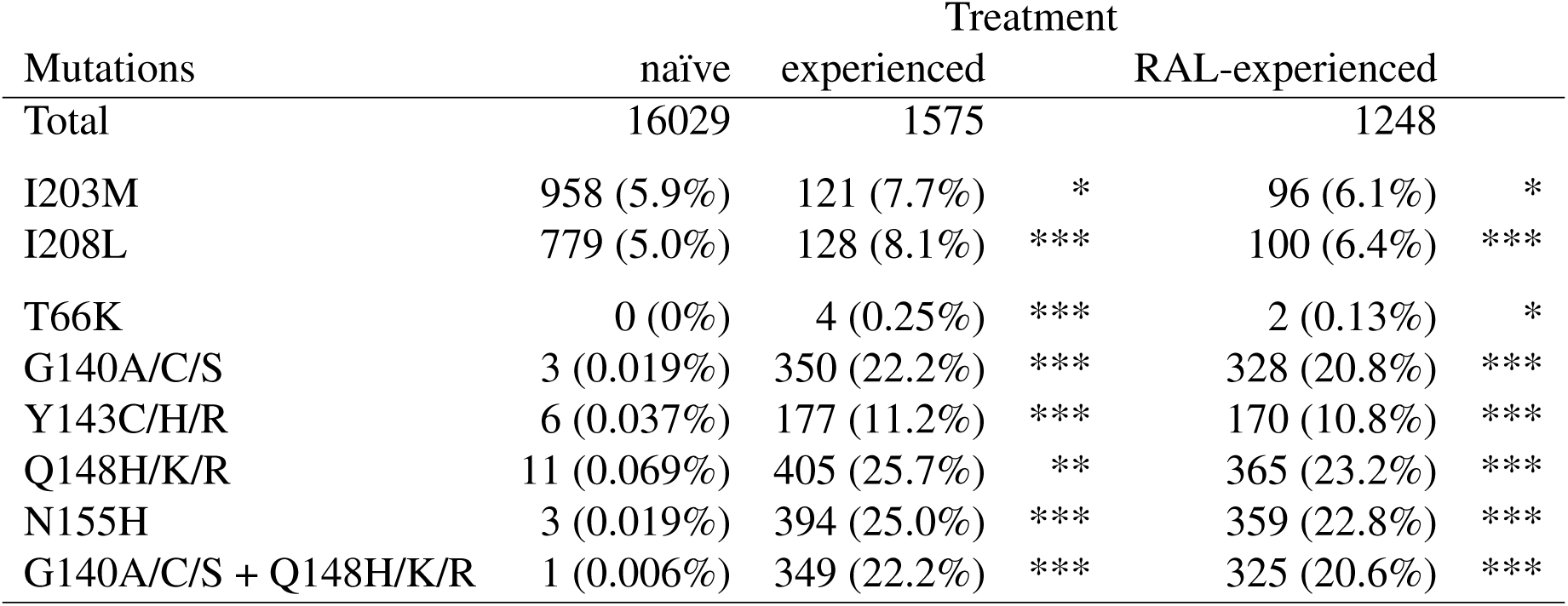
Frequencies of I203M, I208L and RAL INSTI major mutations in HIV-1 IN stratified by treatment according to Stanford HIV Drug Resistance Database (as of February 21, 2019), whereas reported RAL-experienced individuals breakdown is reported. T66K coverage was 16,027 for naïve, I203M and I208L coverage was 15,997 for naïve and 1573 for experienced patients. Other major combinations include: G140A/C/S + Y143C/H/R (9 INI), G140A/C/S + N155H (19 INI), G140A/C/S + Y143C/H/R + Q148H/K/R (9 INI), G140A/C/S + Q148H/K/R + N155H (21 INI), G140A/C/S + Y143C/H/R + Q148H/K/R + N155H (2 INI), Y143C/H/R + Q148H/K/R (10 INI), Y143C/H/R + N155H (26 INI), Y143C/H/R + Q148H/K/R + N155H (2 INI), Q148H/K/R + N155H (27 INI), T66K + G140A/C/S + Q148H/K/R (1 INI), INI = INSTI-experienced, RAL = Raltegravir. Fisher’s exact tests (experienced vs. naïve, RAL- vs. non-RAL-experienced): * = *P <* 0.05; ** = *P <* 0.01; *** = *P <* 0.001.

### Structural analysis

Based on their location in the primary sequence, I203M and I208L maps near the C-terminal base of the *α*-helix connecting the C-terminal domain (CTD) to the catalytic core domain (CCD) of HIV-1 integrase in the unliganded structure [64]. Figure 5A displays the structure of HIV-1 integrase complexed with viral and host DNA (PDB ID 5U1C) and in which only the I203M position could be mapped. However, as discussed in the methods section, a model of HIV-1 integrase was constructed to create an extension of this structure (5U1C extended), such that when superimposed on the original structure (PDB ID 5U1C), the I208L position was roughly mapped.

**Figure 5:**
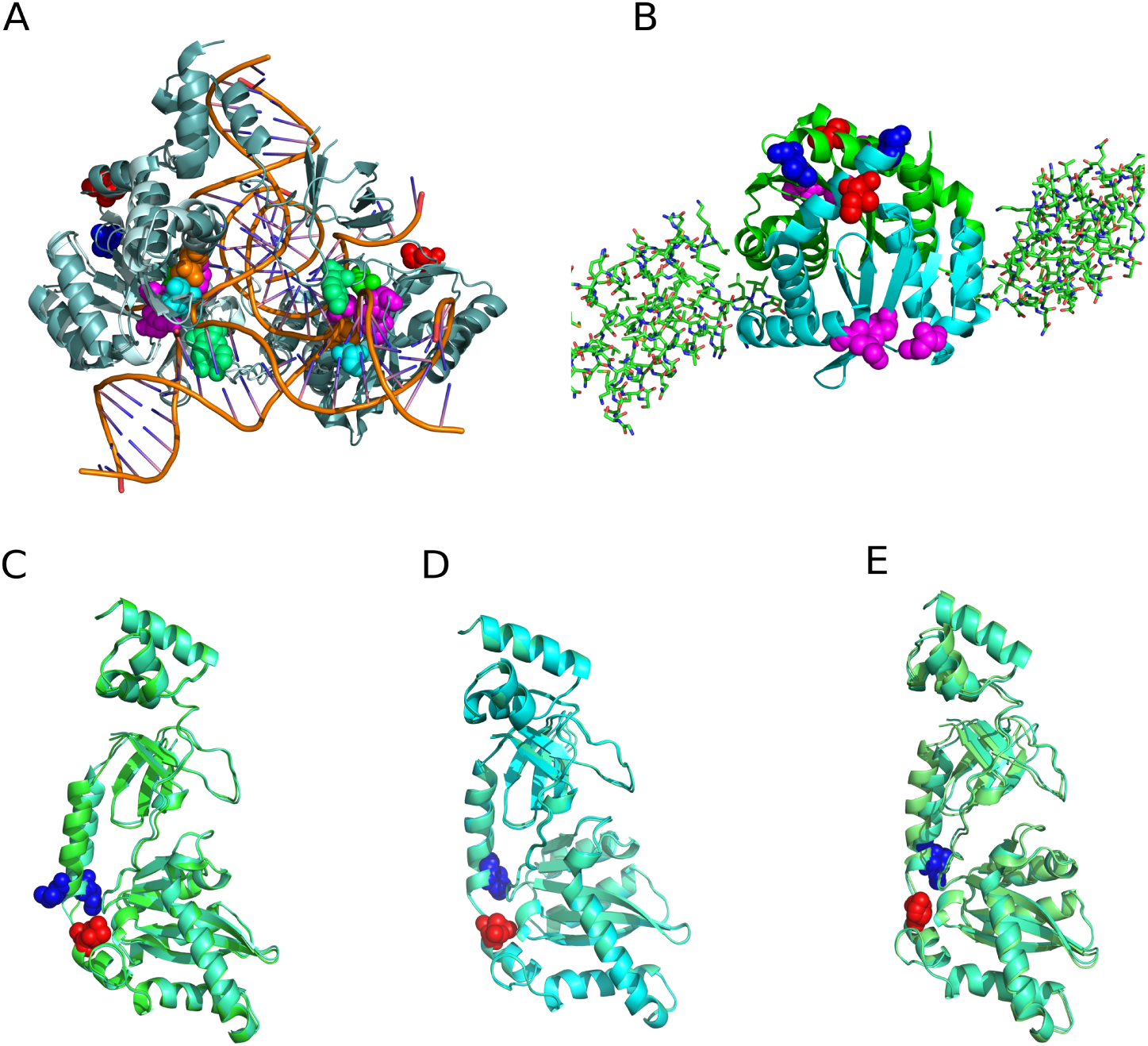
Structural mapping of the novel amino acid replacements. (A) a cryo-electron microscope structure (PDB ID 5U1C) of integrase protein (lightteal) in complex with host and viral DNA. The modelled extended structure (palecyan) is superimposed on chain A to show the missing *α*-helix where the I208L mutation is located (B) a crystal structure of integrase protein CCD (shown as a green cartoon) bound to the human LEDGEF protein (shown as sticks) (C) the modelled I203M mutant structure (limegreen) of HIV-1 integrase superimposed on the modelled extended wild type structure (greencyan) (D) the modelled I208L mutant structure (aquamarine) of HIV-1 integrase superimposed on the modelled extended wild type structure (greencyan) (E) the modelled I203M and I208M mutant structure (lime) of HIV-1 integrase superimposed on the modelled extended wild type structure (greencyan). All catalytic active site residues are colored purple while the novel mutation sites are colored blue (I203M) and red (I208L) respectively. The know major RAL major resistance mutation sites were colored as follows: T66 (cyan), Y143 (lightgreen), Q148 (green), 155 (orange). These images were generated with PyMOL [81].

From both structures, the I203M and I208L residues are distally located from the catalytic active site of HIV DNA processing, the binding site for most INSTI, as well as the location of most drug resistance mutations (all the known major RAL mutations are also shown in the structure in Figure 5A). These residues are closer to the C-terminal domain of HIV-1 integrase that is also responsible for binding to the HIV-1 integration cofactor LEDGF (lens epithelium derived growth factor) and to both host and viral DNA [78–80]. However, structural mapping shows that the I203M and I208L mutations were opposite side of the C-terminal domain engaged in LEDGEF binding (Figure 5B).

Structural modeling of the I203M mutation in HIV-1 integrase shows a conformational change in the *α*-helix connecting the CCD to the CTD (Figure 5C). Similarly the structural model having both the I203M and I208L mutations also show a conformational change the *α*-helix connecting the CCD to the CTD (Figure 5E). The model suggests that these mutations may be conferring INSTI resistance by stabilizing the IN/DNA complex and possibly outlasting the binding of RAL to IN, dependent of the RAL off rate. Based on other drug resistance mechanisms involving differential drug and substrate binding sites, an increase in substrate K_m_/K_off_ (ratio of Michaelis constant to dissociation rate) over inhibitor K_m_/K_off_ can result in resistance. On the other hand, structural modeling of the I208L mutation in HIV-1 integrase shows very little to no conformational changes (Figure 5D). It is however important to stress that these mutations were modeled into a subtype B integrase structure, as structures for other subtypes are not yet available.

## Discussion

The majority of our knowledge on drug resistance mutations (DRMs) in HIV-1 comes from studies of subtype B, even though this subtype represents only a small proportion of infections worldwide. In a previous study [34], we observed an absence of known DRMs associated with RAL resistance in half of the HIV-infected Ugandans failing a third line RAL-based treatment. This absence could be attributed to either complete non-adherence, or a failure to detect resistance-associated polymorphisms below the threshold of detection of Sanger sequencing (about 20% [12]). Based on an initial viral load decrease and adherence tracking during this third line regimen, complete non-adherence resulting in a return of wild type HIV was unlikely for all *n* = 51 patients [34]. Though we cannot retrospectively quantify the extent and impact of drug non-adherence, we can explore the possibility of unique, uncharacterized mutations associated with INSTI resistance with NGS-based genotyping.

Based on previous assay cutoff analyses, the Illumina MiSeq platform is capable of reproducibly detecting mutations at a lower frequency threshold of about 1% [14–16], which confers a substantially improved sensitivity over conventional Sanger sequencing. Setting a frequency threshold of 1% is only meaningful if a sufficient number of virus genomes from the plasma sample are represented in the sequencing library. For instance, if the input number of templates is fewer than 100, then any variant detected at a frequency of 1% or less is likely the result of sequencing error. Our study population comprised patients either sampled prior to initiating ART or following treatment failure. In previous work, we have reported that plasma viral loads averaged about 5.4 log_10_ copies/mL at baseline [8]. Although drug resistance testing in Uganda is requested for patients failing treatment above 1,000 copies/mL, the majority of requests were obtained when viral loads exceed 10,000 copies/mL (averaging 4.8 log_10_ copies/mL for first-line treatment failure, and *>*5.0 log_10_ copies/mL for RAL-based/third-line treatment failure) [31, 34].

Our most conservative estimates are that about one-sixth of the viral RNA from 200*μ*L of plasma was transferred from the sample extraction to the RT-PCR reaction mixture, and that about one-half was converted to cDNA. Given that half of the reaction mixture was used for PCR amplification and sequencing, we would expect at least 850 templates on average to be available for NGS. To evaluate the effect of template resampling on our ability to measure variant frequencies, we can simulate the sampling process assuming that extraction and aliquoting is sampling uniformly at random without replacement, and that sequencing post-amplification is sampling uniformly at random with replacement. For instance, we predict that a variant found in 500 copies/mL (0.5% of the plasma sample) has a 0.03% probability of being sampled to a frequency of 1% or greater under our experimental conditions (2.5% and 97.5% quantiles = 0.26%, 0.77%; *N* = 10^6^ replicates). Conversely, a variant in 1500 copies/mL (1.5%) would be sampled at 1% or less with probability 0.93%. We further note that our multiple imputation across repeated NGS of the same samples would have averaged out some sampling variation.

Our NGS analysis of RAL naïve and treatment failure samples confirms results from our previous study [34], including the absence of previously identified DRMs in about half of RAL failure cases. The SVM analysis of these data also identified a number of potentially novel mutations associated with reduced sensitivity to RAL. Because this classifier evaluates features by selecting data points (the support vectors) to anchor the hyperplane separating labels, some of these features may be associated by chance with RAL failures due to their linkage to features with direct effects. Consequently, we carried out a *post hoc* odds ratio analysis to evaluate the significance of univariate associations between each feature selected by the SVM and the labels (virologic control versus RAL failure). The combined analyses recovered several known major and accessory mutations conferring resistance to RAL. For instance, we found a highly significant association between the major mutation N155H and patients failing RAL treatment [71], and this mutation was significantly linked with accessory mutations such as V151I [74]. A limitation of our SVM analysis was that we selected two frequency thresholds to dichotomize amino acid polymorphisms into binary variables. Although it is possible to directly apply the SVM classifier to continuous variables, dichotomizing the observed frequencies into presence/absence states was necessary to make the analysis computationally feasible. We selected the two frequency thresholds (1% and 20%) to span the range bounded by the lower limits of detection for the Illumina MiSeq [17] and Sanger sequencing [12], respectively. Additionally, the selection of 1% for dichotomizing HIV-1 deep sequence data has also been empirically validated in a recent whole-genome deep sequencing study of HIV-1 [82] and employed in a genome-wide association study of HIV drug resistance [83].

Our SVM analyses identified a cluster of amino acid polymorphisms associated with RAL failure in the *α* helix domain, including the mutations I203M and I208L. With this new information, we could map the majority of RAL failure samples to four largely independent mutational pathways characterized respectively by I203M, I208L, Q148/R, and N155H (Figure 3). We subsequently confirmed these candidate DRMs by introducing the I203M and I208L mutations into a wildtype background through site-directed mutagenesis, and then evaluated those mutants in INSTI susceptibility assays *in vitro*. Our results indicate that both mutations confer a substantial decrease in susceptibility to RAL (Figure 4). A conspicuous limitation of these experiments was the use of the NL4-3 background, a standard laboratory clone in widespread use that was originally derived from an HIV-1 subtype B isolate. Given the high estimates of EC_50_ from our *in vitro* assays, it seems implausible that previous studies of INSTI resistance in HIV-1 subtype B have not observed these mutations. Hence, it is likely that I203M and I208L induce substantial costs to the replicative fitness of the subtype B virus *in vivo*. Further experimental work in our lab will be directed on evaluating drug susceptibilities and fitness costs of these mutations in non-B subtype backgrounds.

Structural analysis shows that the I203M and I208L mutations are located in the CCD, but in sites closer to the CTD rather than the catalytic active sites where DNA is processed and where the integrase inhibitors bind. A previous study suggested that the integrase CTD is also involved in 3’ DNA processing as well as strand transfer to the target DNA [84]. We speculate that these mutations may have resulted in slight conformational changes in the CTD that facilitates increased DNA binding and improved DNA processing, while hindering the actions of RAL. A less likely but alternative speculation was that these mutations may lead to interactions or conformational changes in the CCD of integrase that increases the affinity for DNA while reducing the ability of RAL to bind.

I203M has previously appeared on lists of INSTI resistance mutations in the earlier literature [85–87]. However, it is not currently recognized in actively-maintained clinical guidelines and algorithms, such as the International Antiviral Society-USA Drug Resistance Mutations list [6] or the Stanford HIV Drug Resistance database [23]. Furthermore, I203M was previously characterized as a minor or accessory mutation with no evidence of directly reducing susceptibility, and has also been characterized as a natural polymorphism that is observed a substantial frequencies in untreated individuals [77]. We surmise that the over-representation of subtype B in earlier studies of HIV-1 integrase variation may have obscured the major effect of this mutation on INSTI resistance that we have observed in both machine learning and *in vitro* analyses. In contrast, we have found almost no previous mention of I208L in the HIV-1 drug resistance literature.

Another limitation of our study is that our NGS experiments focused specifically on the region of the HIV-1 genome encoding integrase. For example, recent work has identified mutations outside of this region that can confer resistance to INSTIs, such as a cluster of mutations in the HIV-1 3’ polypurine tract [88], a conserved motif that primes the sythesis of plus-strand HIV-1 DNA during reverse transcription. Newer whole-genome deep sequencing protocols make it possible to apply a similar analytical approach on a global scale, although this will also present both technical and bioinformatic challenges [89]. In particular, expanding coverage will exacerbate the ‘large *p* small *n*’ problem [90] of finding associations for thousands of potential sites with limited sample sizes; *i.e.*, much larger numbers of RAL failures will be needed to support increasing the number of variables by an order of magnitude.

In addition to I203M and I208L, our analysis has identified several other potentially novel RAL resistance mutations in HIV-1 integrase that represent further targets for experimental characterization. Moreover, the cluster of potential mutations spanning IN reference positions 198 to 218, including the mutations I203M and I208L with resistance effects that are independent of the known mutations (Figure 3), implies the existence of a novel and possibly subtype-dependent mechanism of drug resistance that requires further bioinformatic and experimental investigation.

## Acknowledgments

This work was supported in part by Gilead, in part by the Government of Canada through Genome Canada and the Ontario Genomics Institute (OGI-131 to AFYP), by grants from the Canadian Institutes of Health Research (PJT-155990 to AFYP) and partly grants from The National Institute of Allergy and Infectious Diseases/National Institutes of Health (1R56AI139010-01A1 to MEQM) and University of Otago (to MEQM), and in part by Queen Elizabeth II Diamond Jubilee scholarship through Western University DLI O19375892122 that supported EM’s work. We would like to thank Garway T. Ng for assistance with debugging and Dr. Meijuan Tian for lab experiments assistance.

## References

1 Pau AK, George JM. Antiretroviral therapy: current drugs. Infectious Disease Clinics. 2014;28(3):371–402.

2 Wainberg MA, Mesplède T, Quashie PK. The development of novel HIV integrase inhibitors and the problem of drug resistance. Current opinion in virology. 2012;2(5):656–662.

3 Organization WH, et al. Antiretroviral therapy for HIV infection in adults and adolescents: recommendations for a public health approach-2010 revision. 2010;.

4 Fransen S, Gupta S, Danovich R, Hazuda D, Miller M, Witmer M, et al. Loss of raltegravir susceptibility by human immunodeficiency virus type 1 is conferred via multiple nonoverlapping genetic pathways. Journal of virology. 2009;83(22):11440–11446.

5 Llibre JM, Pulido F, García F, Garcia Deltoro M, Blanco JL, Delgado R. Genetic barrier to resistance for dolutegravir. AIDS Rev. 2015;17(1):56–64.

6 Wensing AM, Calvez V, Günthard HF, Johnson VA, Paredes R, Pillay D, et al. 2017 Update of the Drug Resistance Mutations in HIV-1. Topics in antiviral medicine. 2017;24(4):132–133.

7 Liu TF, Shafer RW. Web resources for HIV type 1 genotypic-resistance test interpretation. Clinical infectious diseases. 2006;42(11):1608–1618.

8 Kyeyune F, Gibson RM, Nankya I, Venner C, Metha S, Akao J, et al. Low-frequency drug resistance in HIV-infected Ugandans on antiretroviral treatment is associated with regimen failure. Antimicrobial agents and chemotherapy. 2016;60(6):3380–3397.

9 Gibson RM, Nickel G, Crawford M, Kyeyune F, Venner C, Nankya I, et al. Sensitive detection of HIV-1 resistance to Zidovudine and impact on treatment outcomes in low-to middle-income countries. Infectious diseases of poverty. 2017;6(1):163.

10 Larder B, Kohli A, Kellam P, Kemp S, Kronick M, Henfrey R. Quantitative detection of HIV-1 drug resistance mutations by automated DNA sequencing. Nature. 1993;365(6447):671.

11 Lapointe H, Dong W, Lee G, Bangsberg D, Martin J, Mocello A, et al. HIV drug resistance testing by high-multiplex wide sequencing on the MiSeq instrument. Antimicrobial agents and chemotherapy. 2015;59(11):6824–6833.

12 Schuurman R, Demeter L, Reichelderfer P, Tijnagel J, de Groot T, Boucher C. Worldwide evaluation of DNA sequencing approaches for identification of drug resistance mutations in the human immunodeficiency virus type 1 reverse transcriptase. Journal of clinical microbiology. 1999;37(7):2291–2296.

13 Johnson JA, Li JF, Wei X, Lipscomb J, Irlbeck D, Craig C, et al. Minority HIV-1 drug resistance mutations are present in antiretroviral treatment–naïve populations and associate with reduced treatment efficacy. PLoS medicine. 2008;5(7):e158.

14 Knapp DJ, McGovern RA, Poon AF, Zhong X, Chan D, Swenson LC, et al. Deep sequencing accuracy and reproducibility using Roche/454 technology for inferring co-receptor usage in HIV-1. PloS one. 2014;9(6):e99508.

15 Gibson RM, Meyer AM, Winner D, Archer J, Feyertag F, Ruiz-Mateos E, et al. Sensitive deep-sequencing-based HIV-1 genotyping assay to simultaneously determine susceptibility to protease, reverse transcriptase, integrase, and maturation inhibitors, as well as HIV-1 coreceptor tropism. Antimicrobial agents and chemotherapy. 2014;58(4):2167–2185.

16 Van Laethem K, Theys K, Vandamme AM. HIV-1 genotypic drug resistance testing: digging deep, reaching wide? Current opinion in virology. 2015;14:16–23.

17 Brumme CJ, Poon AF. Promises and pitfalls of Illumina sequencing for HIV resistance genotyping. Virus research. 2017;239:97–105.

18 Pfeifer N, Lengauer T. Improving HIV coreceptor usage prediction in the clinic using hints from next-generation sequencing data. Bioinformatics. 2012;28(18):i589–i595.

19 Olejnik M, Steuwer M, Gorlatch S, Heider D. gCUP: rapid GPU-based HIV-1 co-receptor usage prediction for next-generation sequencing. Bioinformatics. 2014;30(22):3272–3273.

20 Lee GQ, Harrigan PR, Dong W, Poon AF, Heera J, Demarest J, et al. Comparison of population and 454 deep sequence analysis for HIV type 1 tropism versus the original trofile assay in non-B subtypes. AIDS research and human retroviruses. 2013;29(6):979–984.

21 Vallari A, Holzmayer V, Harris B, Yamaguchi J, Ngansop C, Makamche F, et al. Confirmation of putative HIV-1 group P in Cameroon. Journal of virology. 2011;85(3):1403–1407.

22 Tebit DM, Arts EJ. Tracking a century of global expansion and evolution of HIV to drive understanding and to combat disease. The Lancet infectious diseases. 2011;11(1):45–56.

23 Shafer RW. Rationale and uses of a public HIV drug-resistance database. The Journal of infectious diseases. 2006;194(Supplement 1):S51–S58.

24 Hemelaar J, Gouws E, Ghys PD, Osmanov S. Global and regional distribution of HIV-1 genetic subtypes and recombinants in 2004. Aids. 2006;20(16):W13–W23.

25 Bocket L, Cheret A, Deuffic-Burban S, Choisy P, Gerard Y, de la Tribonnière X, et al. Impact of human immunodeficiency virus type 1 subtype on first-line antiretroviral therapy effectiveness. Antivir Ther. 2005;10(2):247–54.

26 Geretti AM, Harrison L, Green H, Sabin C, Hill T, Fearnhill E, et al. Effect of HIV-1 subtype on virologic and immunologic response to starting highly active antiretroviral therapy. Clinical infectious diseases. 2009;48(9):1296–1305.

27 Easterbrook PJ, Smith M, Mullen J, O’Shea S, Chrystie I, de Ruiter A, et al. Impact of HIV-1 viral subtype on disease progression and response to antiretroviral therapy. Journal of the International AIDS Society. 2010;13(1):4.

28 Bhargava M, Cajas JM, Wainberg MA, Klein MB, Pai NP. Do HIV-1 non-B subtypes differentially impact resistance mutations and clinical disease progression in treated populations? Evidence from a systematic review. Journal of the International AIDS Society. 2014;17(1).

29 Brenner B, Turner D, Oliveira M, Moisi D, Detorio M, Carobene M, et al. A V106M mutation in HIV-1 clade C viruses exposed to efavirenz confers cross-resistance to non-nucleoside reverse transcriptase inhibitors. Aids. 2003;17(1):F1–F5.

30 Malet I, Fourati S, Charpentier C, Morand-Joubert L, Armenia D, Wirden M, et al. The HIV-1 integrase G118R mutation confers raltegravir resistance to the CRF02 AG HIV-1 subtype. Journal of Antimicrobial Chemotherapy. 2011;66(12):2827–2830.

31 Poon AF, Ndashimye E, Avino M, Gibson R, Kityo C, Kyeyune F, et al. First-line HIV treatment failures in non-B subtypes and recombinants: a cross-sectional analysis of multiple populations in Uganda. AIDS research and therapy. 2019;16(1):3.

32 Kyeyune F, Nankya I, Metha S, Akao J, Ndashimye E, Tebit DM, et al. Treatment failure and drug resistance is more frequent in HIV-1 subtype D versus subtype A-infected Ugandans over a 10-year study period. AIDS (London, England). 2013;27(12):1899.

33 World Health Organization. Update of recommendations on first- and second-line antiretroviral regimens; 2019. Available from: https://apps.who.int/iris/bitstream/handle/10665/325892/WHO-CDS-HIV-19.15-eng.pdf.

34 Ndashimye E, Avino M, Kyeyune F, Nankya I, Gibson RM, Nabulime E, et al. Absence of HIV-1 drug resistance mutations supports the use of dolutegravir in Uganda. AIDS research and human retroviruses. 2018;34(5):404–414.

35 Günthard HF, Aberg JA, Eron JJ, Hoy JF, Telenti A, Benson CA, et al. Antiretroviral treatment of adult HIV infection: 2014 recommendations of the International Antiviral Society–USA Panel. Jama. 2014;312(4):410–425.

36 Dorward J, Lessells R, Drain PK, Naidoo K, de Oliveira T, Pillay Y, et al. Dolutegravir for first-line antiretroviral therapy in low-income and middle-income countries: uncertainties and opportunities for implementation and research. The Lancet HIV. 2018;5(7):e400–e404.

37 Gatell JM, Katlama C, Grinsztejn B, Eron JJ, Lazzarin A, Vittecoq D, et al. Long-term efficacy and safety of the HIV integrase inhibitor raltegravir in patients with limited treatment options in a Phase II study. JAIDS Journal of Acquired Immune Deficiency Syndromes. 2010;53(4):456–463.

38 Steigbigel RT, Cooper DA, Teppler H, Eron JJ, Gatell JM, Kumar PN, et al. Long-term efficacy and safety of Raltegravir combined with optimized background therapy in treatment-experienced patients with drug-resistant HIV infection: week 96 results of the BENCHMRK 1 and 2 Phase III trials. Clinical Infectious Diseases. 2010;50(4):605–612.

39 Eron JJ, Cooper DA, Steigbigel RT, Clotet B, Gatell JM, Kumar PN, et al. Efficacy and safety of raltegravir for treatment of HIV for 5 years in the BENCHMRK studies: final results of two randomised, placebo-controlled trials. The Lancet Infectious Diseases. 2013;13(7):587–596.

40 Hu Z, Kuritzkes DR. Effect of raltegravir resistance mutations in HIV-1 integrase on viral fitness. Journal of acquired immune deficiency syndromes (1999). 2010;55(2):148.

41 Malet I, Delelis O, Valantin MA, Montes B, Soulie C, Wirden M, et al. Mutations associated with failure of raltegravir treatment affect integrase sensitivity to the inhibitor in vitro. Antimicrobial agents and chemotherapy. 2008;52(4):1351–1358.

42 Delelis O, Thierry S, Subra F, Simon F, Malet I, Alloui C, et al. Impact of Y143 HIV-1 integrase mutations on resistance to raltegravir in vitro and in vivo. Antimicrobial agents and chemotherapy. 2010;54(1):491–501.

43 Björndal A, Deng H, Jansson M, Fiore JR, Colognesi C, Karlsson A, et al. Coreceptor usage of primary human immunodeficiency virus type 1 isolates varies according to biological phenotype. Journal of virology. 1997;71(10):7478–7487.

44 Dudley DM, Gao Y, Nelson KN, Henry KR, Nankya I, Gibson RM, et al. A novel yeast-based recombination method to clone and propagate diverse HIV-1 isolates. Biotechniques. 2009;46(6):458–467.

45 Kimpton J, Emerman M. Detection of replication-competent and pseudotyped human immunodeficiency virus with a sensitive cell line on the basis of activation of an integrated beta-galactosidase gene. Journal of virology. 1992;66(4):2232–2239.

46 Martin M. Cutadapt removes adapter sequences from high-throughput sequencing reads. EMBnet journal. 2011;17(1):10–12.

47 Langmead B, Salzberg SL. Fast gapped-read alignment with Bowtie 2. Nature methods. 2012;9(4):357.

48 Aguas R, Ferguson NM. Feature selection methods for identifying genetic determinants of host species in RNA viruses. PLoS computational biology. 2013;9(10):e1003254.

49 García-Laencina PJ, Sancho-Gómez JL, Figueiras-Vidal AR. Pattern classification with missing data: a review. Neural Computing and Applications. 2010;19(2):263–282.

50 Buuren S, Groothuis-Oudshoorn K. mice: Multivariate imputation by chained equations in R. Journal of statistical software. 2011;45(3).

51 White IR, Royston P, Wood AM. Multiple imputation using chained equations: issues and guidance for practice. Statistics in medicine. 2011;30(4):377–399.

52 Srebro N, Rennie J, Jaakkola TS. Maximum-margin matrix factorization. In: Advances in neural information processing systems; 2005. p. 1329–1336.

53 Becker N, Werft W, Toedt G, Lichter P, Benner A. penalizedSVM: a R-package for feature selection SVM classification. Bioinformatics. 2009;25(13):1711–1712.

54 Bradley PS, Mangasarian OL. Feature selection via concave minimization and support vector machines. In: ICML. vol. 98; 1998. p. 82–90.

55 Becker N, Toedt G, Lichter P, Benner A. Elastic SCAD as a novel penalization method for SVM classification tasks in high-dimensional data. BMC bioinformatics. 2011;12(1):138.

56 Pagano M, Gauvreau K. Principles of biostatistics. 2000. Brooks/Cole: India;2.

57 Pond SLK, Posada D, Stawiski E, Chappey C, Poon AF, Hughes G, et al. An evolutionary model-based algorithm for accurate phylogenetic breakpoint mapping and subtype prediction in HIV-1. PLoS computational biology. 2009;5(11):e1000581.

58 Edgar RC. MUSCLE: multiple sequence alignment with high accuracy and high throughput. Nucleic acids research. 2004;32(5):1792–1797.

59 Larsson A. AliView: a fast and lightweight alignment viewer and editor for large datasets. Bioinformatics. 2014;30(22):3276–3278.

60 Darriba D, Taboada GL, Doallo R, Posada D. jModelTest 2: more models, new heuristics and parallel computing. Nature methods. 2012;9(8):772–772.

61 Guindon S, Dufayard JF, Lefort V, Anisimova M, Hordijk W, Gascuel O. New algorithms and methods to estimate maximum-likelihood phylogenies: assessing the performance of PhyML 3.0. Systematic biology. 2010;59(3):307–321.

62 Šali A, Blundell TL. Comparative protein modelling by satisfaction of spatial restraints. Journal of molecular biology. 1993;234(3):779–815.

63 Bhattacharya D, Nowotny J, Cao R, Cheng J. 3Drefine: an interactive web server for efficient protein structure refinement. Nucleic acids research. 2016;44(W1):W406–W409.

64 Chen JCH, Krucinski J, Miercke LJ, Finer-Moore JS, Tang AH, Leavitt AD, et al. Crystal structure of the HIV-1 integrase catalytic core and C-terminal domains: a model for viral DNA binding. Proceedings of the National Academy of Sciences. 2000;97(15):8233–8238.

65 Schirmer M, DAmore R, Ijaz UZ, Hall N, Quince C. Illumina error profiles: resolving fine-scale variation in metagenomic sequencing data. BMC bioinformatics. 2016;17(1):125.

66 Cristiani N, Shawe-Taylor J. An introduction to support vector machines and other kernel-based learning methods. Cambridge University Press; 2000.

67 Cai YD, Liu XJ, Xu XB, Chou KC. Support vector machines for predicting HIV protease cleavage sites in protein. Journal of Computational Chemistry. 2002;23(2):267–274.

68 Darnag R, Mazouz EM, Schmitzer A, Villemin D, Jarid A, Cherqaoui D. Support vector machines: development of QSAR models for predicting anti-HIV-1 activity of TIBO derivatives. European journal of medicinal chemistry. 2010;45(4):1590–1597.

69 Margot NA, Hluhanich RM, Jones GS, Andreatta KN, Tsiang M, McColl DJ, et al. In vitro resistance selections using elvitegravir, raltegravir, and two metabolites of elvitegravir M1 and M4. Antiviral research. 2012;93(2):288–296.

70 Goethals O, Clayton R, Van Ginderen M, Vereycken I, Wagemans E, Geluykens P, et al. Resistance mutations in human immunodeficiency virus type 1 integrase selected with elvitegravir confer reduced susceptibility to a wide range of integrase inhibitors. Journal of virology. 2008;82(21):10366–10374.

71 Fransen S, Gupta S, Frantzell A, Petropoulos CJ, Huang W. Substitutions at amino acid positions 143, 148, and 155 of HIV-1 integrase define distinct genetic barriers to raltegravir resistance in vivo. Journal of virology. 2012;86(13):7249–7255.

72 Van Wesenbeeck L, Rondelez E, Feyaerts M, Verheyen A, Van der Borght K, Smits V, et al. Cross-resistance profile determination of two second-generation HIV-1 integrase inhibitors using a panel of recombinant viruses derived from raltegravir-treated clinical isolates. Antimicrobial agents and chemotherapy. 2011;55(1):321–325.

73 Kobayashi M, Nakahara K, Seki T, Miki S, Kawauchi S, Suyama A, et al. Selection of diverse and clinically relevant integrase inhibitor-resistant human immunodeficiency virus type 1 mutants. Antiviral research. 2008;80(2):213–222.

74 Blanco JL, Varghese V, Rhee SY, Gatell JM, Shafer RW. HIV-1 integrase inhibitor resistance and its clinical implications. Journal of Infectious Diseases. 2011;203(9):1204–1214.

75 Rhee SY, Sankaran K, Varghese V, Winters MA, Hurt CB, Eron JJ, et al. HIV-1 protease, reverse transcriptase, and integrase variation. Journal of virology. 2016;90(13):6058–6070.

76 Goethals O, Vos A, Van Ginderen M, Geluykens P, Smits V, Schols D, et al. Primary mutations selected in vitro with raltegravir confer large fold changes in susceptibility to first-generation integrase inhibitors, but minor fold changes to inhibitors with second-generation resistance profiles. Virology. 2010;402(2):338–346.

77 Rhee SY, Liu TF, Kiuchi M, Zioni R, Gifford RJ, Holmes SP, et al. Natural variation of HIV-1 group M integrase: implications for a new class of antiretroviral inhibitors. Retrovirology. 2008;5(1):74.

78 Heuer TS, Brown PO. Mapping Features of HIV-1 Integrase Near Selected Sites on Viral and Target DNA Molecules in an Active Enzyme-DNA Complex by Photo-Cross-Linking. Biochemistry. 1997;36(35):10655–10665.

79 Kessl J, McKee C, Eidahl J, Shkriabai N, Katz A, Kvaratskhelia M. HIV-1 integrase-DNA recognition mechanisms. Viruses. 2009;1(3):713–736.

80 Roberts VA. C-terminal domain of integrase binds between the two active sites. Journal of chemical theory and computation. 2015;11(9):4500–4511.

81 DeLano WL. Pymol: An open-source molecular graphics tool. CCP4 Newsletter On Protein Crystallography. 2002;40(1):82–92.

82 Zanini F, Brodin J, Thebo L, Lanz C, Bratt G, Albert J, et al. Population genomics of intrapatient HIV-1 evolution. Elife. 2015;4:e11282.

83 Power RA, Davaniah S, Derache A, Wilkinson E, Tanser F, Gupta RK, et al. Genome-Wide Association Study of HIV Whole Genome Sequences Validated using Drug Resistance. PLOS ONE. 2016 09;11:1–14. Available from: https://doi.org/10.1371/journal.pone.0163746.

84 Chiu TK, Davies DR. Structure and function of HIV-1 integrase. Current topics in medicinal chemistry. 2004;4(9):965–977.

85 Lataillade M, Chiarella J, Kozal MJ. Short communication Natural polymorphism of the HIV-1 integrase gene and mutations associated with integrase inhibitor resistance. Antiviral therapy. 2007;12:563–570.

86 Garrido C, Geretti AM, Zahonero N, Booth C, Strang A, Soriano V, et al. Integrase variability and susceptibility to HIV integrase inhibitors: impact of subtypes, antiretroviral experience and duration of HIV infection. Journal of antimicrobial chemotherapy. 2009;65(2):320–326.

87 Da Silva D, Van Wesenbeeck L, Breilh D, Reigadas S, Anies G, Van Baelen K, et al. HIV-1 resistance patterns to integrase inhibitors in antiretroviral-experienced patients with viro-logical failure on raltegravir-containing regimens. Journal of Antimicrobial Chemotherapy. 2010;65(6):1262–1269.

88 Malet I, Subra F, Charpentier C, Collin G, Descamps D, Calvez V, et al. Mutations located outside the integrase gene can confer resistance to HIV-1 integrase strand transfer inhibitors. MBio. 2017;8(5):e00922–17.

89 Zanini F, Brodin J, Albert J, Neher RA. Error rates, PCR recombination, and sampling depth in HIV-1 whole genome deep sequencing. Virus research. 2017;239:106–114.

90 Dutilh BE, Backus L, Edwards RA, Wels M, Bayjanov JR, van Hijum SA. Explaining microbial phenotypes on a genomic scale: GWAS for microbes. Briefings in functional genomics. 2013;12(4):366–380.

